# Regulation associated modules reflect 3D genome modularity associated with chromatin activity

**DOI:** 10.1101/2022.03.02.482718

**Authors:** Lina Zheng, Wei Wang

## Abstract

The 3D genome has been shown to be organized into modules including topologically associating domains (TADs) and compartments that are primarily defined by spatial contacts from Hi-C or other experiments. There exists a gap to investigate whether and how the spatial modularity of the chromatin is related to the functional modularity resulting from the chromatin activity. Increasing evidence shows a tight interplay between histone modifications and 3D chromatin organization. As the histone modifications reflect the chromatin activity, it is tempting to infer the spatial modularity of the genome directly from the histone modification patterns, which would establish the connection between the spatial and functional modularity of the genome. However, uncovering the 3D genomic modules using histone modifications has not been well explored. Here, we report that the histone modifications show a modular pattern (referred to as regulation associated modules, RAMs) that reflects the spatial modularity of the chromatin structure. We found that enhancer-promoter interactions and extrachromosomal DNAs (ecDNAs) occur more often within the same RAMs than within the same TADs, indicating stronger insulation of the RAM boundaries and a modularization of the 3D genome at a scale better aligned with the chromatin activity. Consistently, compared to the TAD boundaries, in silico predictions showed that deletions of RAM boundaries perturb the chromatin structure more severely and somatic variants in the cancer samples are more enriched in the RAM boundaries. These observations suggest that RAMs reflect a modular organization of the 3D genome at a scale better aligned with chromatin activity, providing a bridge connecting the structural and functional modularity of the genome.

## INTRODUCTION

Histone modifications are critical to shape the chromatin structure and regulate gene expression(*1*, *2*). Active marks such as H3K27ac and H3K4me3 open up chromatin to allow access of transcription factors (TFs) and transcription machinery to promoters or enhancers. Repressive marks such as H3K9me3 and H3K27me3 condensate chromatin to block TF binding and suppress gene expression. DNA marked by active and repressive histone modifications form euchromatin and heterochromatin that are distinct on the compactness. These observations suggest that histone modifications have an important impact on organizing the regional and global 3D genome.

Accumulating evidence has revealed the association of histone modifications with the topologically associating domains (TADs)(*3*–*6*) and compartments(*7*, *8*) derived from the Hi-C contact maps showing plaid patterns. TADs represent genomic domains forming dense internal contacts but fewer contacts with neighboring regions. The TAD boundaries are demarcated with CTCF sites or active transcribed DNA sequences. The Hi-C data also shows that the 3D genome is partitioned into transcriptionally active (compartment A) and suppressed (compartment B) compartments. Active and repressive histone marks are enriched, but do not exclusively appear, in the A and B compartments, respectively (*7*, *8*). Computational models have shown that histone modification signals are predictive of Hi-C contacts particularly for enhancer-promoter interactions(*9*), TAD boundaries(*10*) and compartments(*11*). Histone modifications are tightly associated with transcriptional activity (*12*–*14*) while transcription and proteins involved in transcriptional regulation including RNA polymerase and TFs have been shown to contribute to compartmentation and active promoters and enhancers tend to form clusters in the nucleus(*15*–*18*).

Despite the mechanisms underlying the interplay between histone modifications and chromatin organization remain elusive, histone modifications can indicate the spatial organization of the genome as readout signals for regulatory modules. However, the current analysis has been limited to associating histone marks to Hi-C derived TADs and compartments. An unfilled gap is to use histone modifications to directly elucidate the modular organization of the 3D genome. We propose here to define the spatial module of the genome organization resulting from the chromatin activities reflected by histone modifications.

We found that the frequency profiles of the H3K27ac peaks present a modular structure (referred to as regulation associated modules, RAMs). A large number of these modules are shared across cell types and can be independently derived using other active histone marks, including H3K4me3 and H3K4me1. We uncovered several lines of evidence to support the hypothesis that the RAMs are spatial modules resulting from functional activities: the enhancer-promoter interactions dominantly occur within RAMs; the extrachromosomal DNAs (ecDNAs) tend to be originated from the same RAMs rather than split in multiple RAMs; RAMs are resistant to cohesin degradation. These properties of RAMs distinguish them from TADs and compartments. Furthermore, deletion of the RAM boundaries is predicted to alter the chromatin organization more significantly than the deletion of TAD boundaries. Consistently, the somatic genetic variations in cancer patients are enriched in RAM boundaries, suggesting a possible mechanism of tumorigenesis involved in altering the chromatin modules.

## RESULTS

### Regulation associated modules (RAMs) detected by the histone modification peaks

We analyzed the density profile of H3K27ac peaks (i.e. peak count in a sliding window) from chromatin immunoprecipitation assays with sequencing (ChIP-Seq) experiments as using the peak density instead of read count density can better remove the noise from the background signals. We downloaded ChIP-seq data of 93 normal and 19 cancer samples from Roadmap Epigenomics Project (http://www.roadmapepigenomics.org/)(*19*) and ENCODE portal (https://www.encodeproject.org/)(*20*) (**Table S1**). Using a sliding window (a fixed flanking size of 500kbp and step size varying from 10kbp to 500kbp), we computed the H3K27ac peak densities in the linear genome. Regardless of the step size, the H3K27ac peak densities were not evenly distributed and showed a modular pattern (**Figure 1A**). The active marks of H3K4me1, H3K4me3 and H3K36me3 showed similar peak density profiles to H3K27ac in the 93 samples, indicated by high Pearson correlations between them, whereas the repressive marks of H3K27me3 and H3K9me3 had less consistent patterns (**Figure 1B**). Given the highly correlated active mark patterns, we focused on analyzing the H3K27ac signals as the other active marks show similar modular structure.

**Figure 1.**
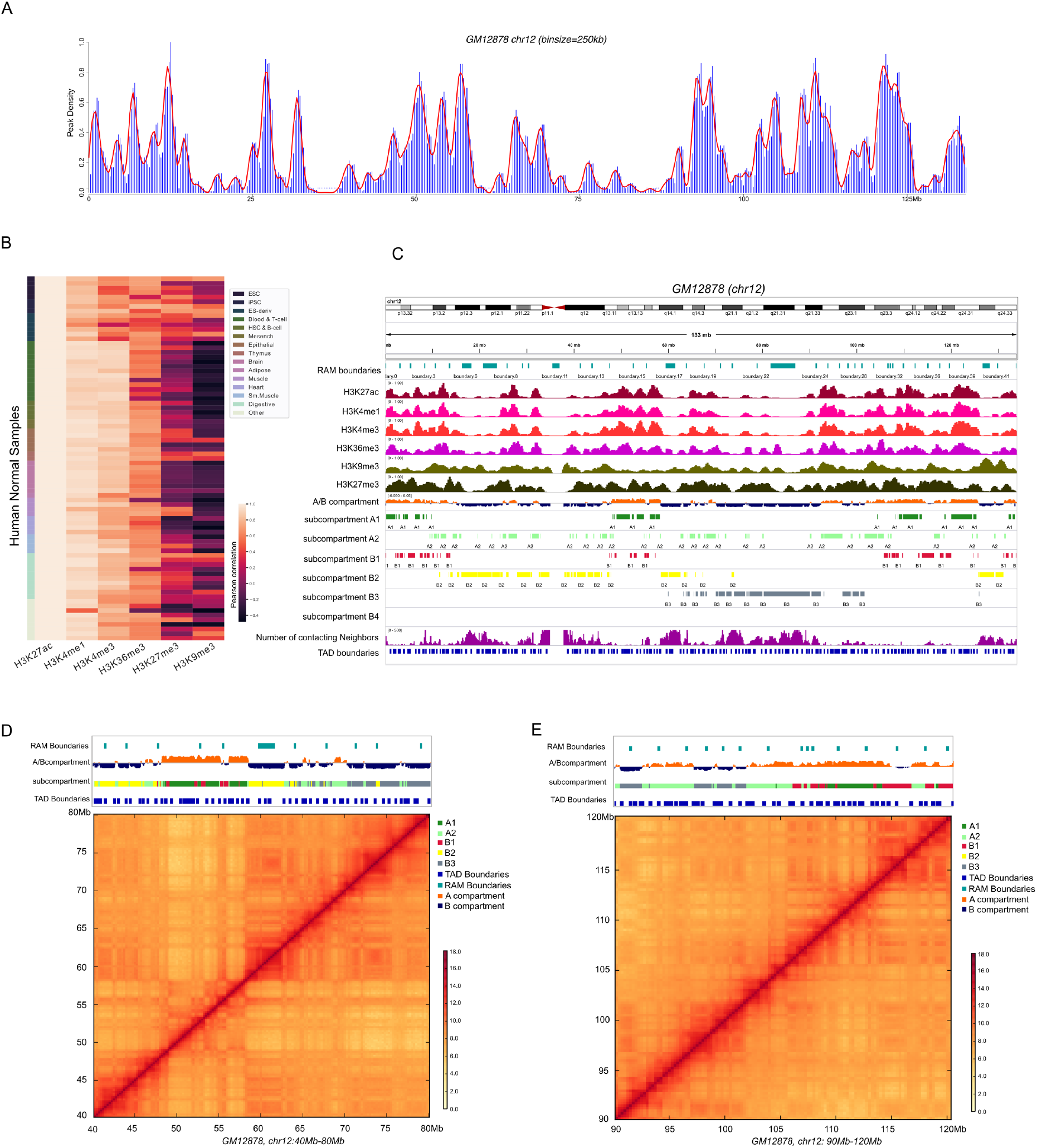
Regulation associated module (RAM) identification. **(A)** H3K27ac peaks density of chr12 in GM12878 (binsize=250kb) **(B)** Pearson correlation between histone modification marks (H3K27ac, H3K4me1, H3K4me3, H3K36me3, H3K27me3, and H3K9me3) for the Roadmap samples. **(C)** Examples of histone modifications, A/B compartments, subcompartments, number of the 3D contacts, TAD boundaries and RAM boundaries in chr12 for GM12878. **(D)** The zoom-in genomic view for chr12:40Mb-80Mb and **(E)** The zoom-in genomic view for chr12:90Mb-120Mb in hg19.

At a given step size, we identified the valley or minima of the H3K27ac peak profile that was smoothed using local polynomial fit in each chromosome and in each cell type (see **Materials and Methods**). These valleys demarcated the boundaries of the modular domains (called Regulation Associated Domains or RAMs). We varied the step size from 10kbp to 500kbp and fixed the window size to 500kbp. It is not surprising that, with the increasing step size, the RAM size increased and a higher percentage of RAMs were shared between samples (**fig S1-S2**). We observed that the number of common RAMs in all the chromosomes reached a plateau at 250kbp step size in both normal and cancer samples, which indicates the identified RAMs are most conserved across diverse cell types (**fig S3A-B**). We thus used this step size of 250kbp for the remaining analyses. A RAM boundary is called consensus RAM (cRAM) boundary if it is shared by >25% of the samples. This way, 711 cRAMs were detected in the normal samples and 771 cRAMs in the cancer samples (see **Materials and Methods**). On average, 60% of the RAMs in a cell type are consensus (referred to as cRAMs) and the remaining cell-type specific (**fig S3C**).

One example of the identified RAMs in chr12 of the GM12878 cell by IGV software(*21*) is shown in **Figure 1C-E**. Obviously, the RAM boundaries have lower signals of the active histone marks (H3K27ac, H3K4me1, H3K4me3, H3K36me3) and higher repressive marks (H3K9me3 and H3K27me3) compared to the within RAM regions. Consistently, they tend to align with the B compartment or subcompartments (B1, B2, B3). Furthermore, by counting the number of the 3D contacting neighbors for each locus using the 10kb resolution Hi-C data in GM12878 (contacts with log(P-value) <=−10), we found that the RAM boundaries tend to harbor many 3D contacts, indicated an enrichment with densely packed DNA sequences forming many spatial contacts (**Figure 1C**).

### Characterization of the consensus regulation associated modules (cRAMs)

As cRAMs are largely shared between diverse cell lines, we further characterized them. Among the cell lines that have both active and repressive marks (all are normal cells), as expected, we found that the cRAM boundaries have lower peak density of the active marks (H3K27ac, H3K4me3, H3K4me1 and H3K36me3) and slightly higher peak densities of repressive marks (H3K27me3 and H3K9me3) than the cRAM regions (see examples in GM12878 and HUVEC cell lines in **Figure 2A**). To quantify the difference of the histone modifications among the cRAM boundaries and the non-boundary regions, we counted the peak density using a sliding window, and compared the histone modifications enrichment across 93 normal samples. The P-value < 0.05 from the Wilcoxon Rank Sum test indicated that cRAM boundaries have significantly lower active marks and higher repressive marks than the non-boundaries of cRAMs (**Figure 2B-G**). Furthermore, using the available 10kb-resolution Hi-C data in the K562, GM12878, A549, IMR90, NHEK, HUVEC, HMEC and HCT116 cell lines, we found that the cRAM boundaries have significantly more Hi-C contacts (intrachromosomal contacts with log(P-value)<=−10) compared to the whole genome (**Figure 2H**), which is consistent with the genome browser view for any RAM boundary in **Figure 1C**. These observations suggested that the cRAM boundaries are formed by densely packed DNA sequences harboring many 3D contacts.

**Figure 2.**
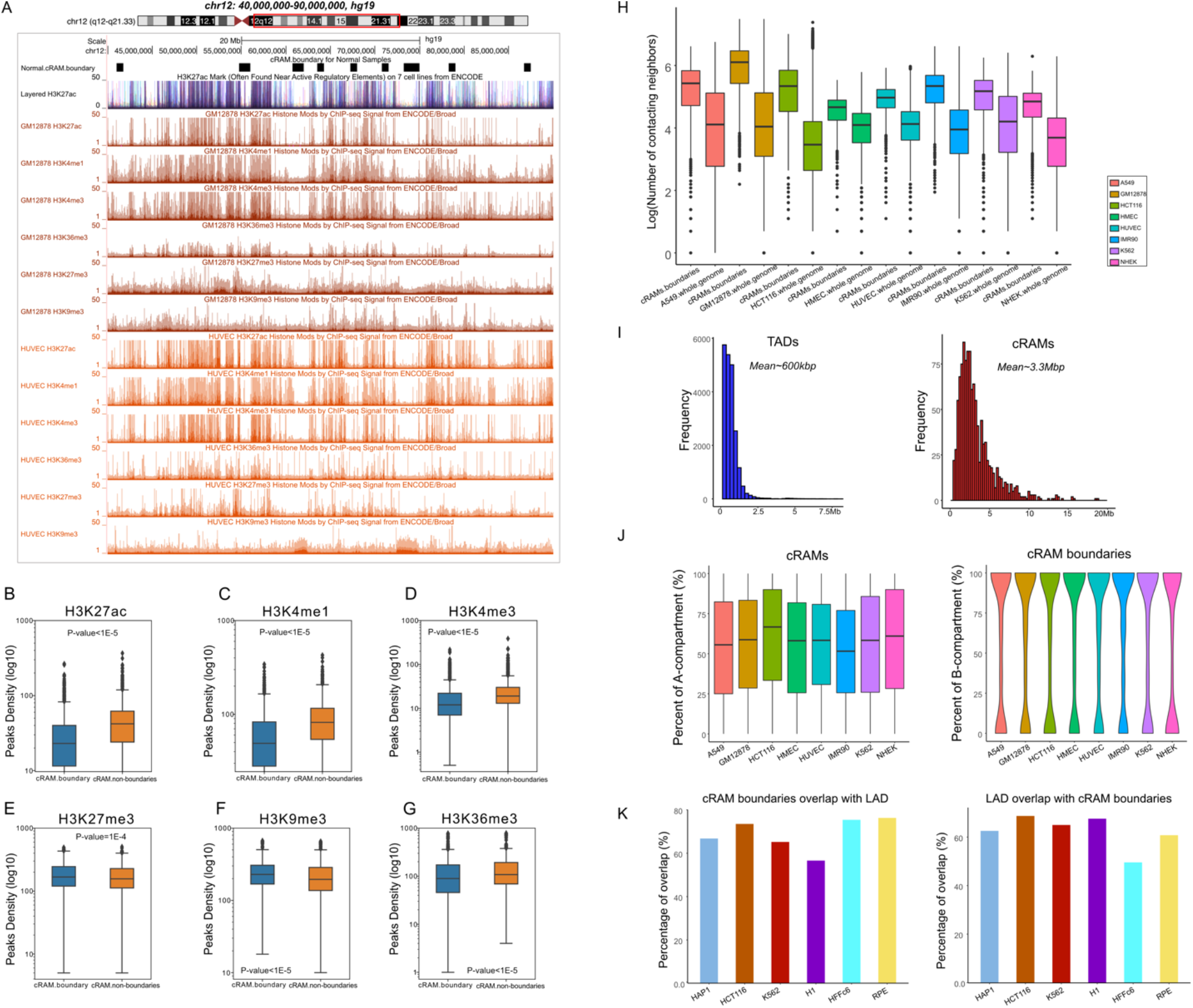
Characterization of the cRAMs and boundaries. (**A**) Genome browser examples of normal cRAM boundaries and the histone modifications in genomic region chr12:40Mb-90Mb in hg19 for GM12878 and the HUVEC cells. (**B-G**) The genome-wide enrichment of the histone modification marks. (**B**) H3K27ac, (**C**) H3K4me1, (**D**) H3K4me3, (**E**) H3K27me3, (**F**) H3K9me3, and (**G**) H3K36me3 in cRAM boundaries and non-boundaries (**H**) The contacting neighbors distribution of cRAM boundaries and whole genome locus in the 3D contact network in a diverse of the cell types. (**I**) Sizes of the TADs and cRAMs. (**J**) cRAM boundaries distribution over A/B compartments. (**K**) cRAM boundaries distribution over LaminB1 signals (LAD).

We next investigated how RAMs are related to the previously identified chromatin modules. First, the median size of cRAMs (~3.3Mbp) is larger than TADs (~600kbp) and one RAM often spans across multiple TADs (**Figure 2I**). Second, using the Hi-C data, we identified the A/B compartments at 250kb resolution (see **Materials and Methods**). We calculated the percentage of the A and B compartments in each cRAM (250kb bin size) across the cell types. While the A-compartments account around 50%-75% in each of the cRAM, a single cRAM is largely composed by a mixture of A and B compartments, indicating a distinction between cRAMs and compartments (**Figure 2J**). Consistently, the cRAM boundaries are enriched with B-compartment but also with a significant portion of A compartments (**Figure 2J**). Third, we checked the Lamin-B1 signals for the cRAM boundaries. Lamin-B1 is a scaffolding component of the nuclear envelope(*22*, *23*). A positive signal for Lamin-B1 suggests a close distance to the nuclear lamina, which could be used to define lamina associated domain (LAD). When aligning the cRAM boundaries with the Lamin-B1 signals (see **Materials and Methods**), we found on average 69% of the cRAM boundaries overlapping with Lamin-B1 signals across the cell types and meanwhile on average 62.7% of the LADs identified from each cell type overlapping with cRAM boundaries (**Figure 2K**), indicating that LADs and cRAMs are also different. Taken together, the cRAM boundaries are formed by densely packed DNA sequences; while they are enriched with B compartment and Lamin-B1 signals, RAMs are clearly distinct from the previously reported domain structures such as TADs, LADs and A/B compartments.

### RAMs are functional units

If RAMs are functional modules, we reason that the majority of the promoter-enhancer interactions should occur within the same RAMs. We downloaded 970 high-confidence promoter-enhancer interactions in the K562 cell line that were experimentally validated in ref (*24*, *25*) and 95% of them are located within the same RAMs, compared to 75% of them in the same TADs(*4*, *6*, *8*, *26*) (**Figure 3A-B**). Two examples of promoter-enhancer interactions are shown in **Figure 3G**: the enhancer-promoter interactions of *STEAP1B* and *VGF* are across multiple TADs but within the same K562 RAMs marked by continuous strong H3K27ac peaks. This observation suggests that RAMs may represent regulatory modules and RAM boundaries insulate promoter-enhancer contacts across RAMs at a scale more appropriate than TADs to capture functional modularity of chromatin activity.

**Figure 3.**
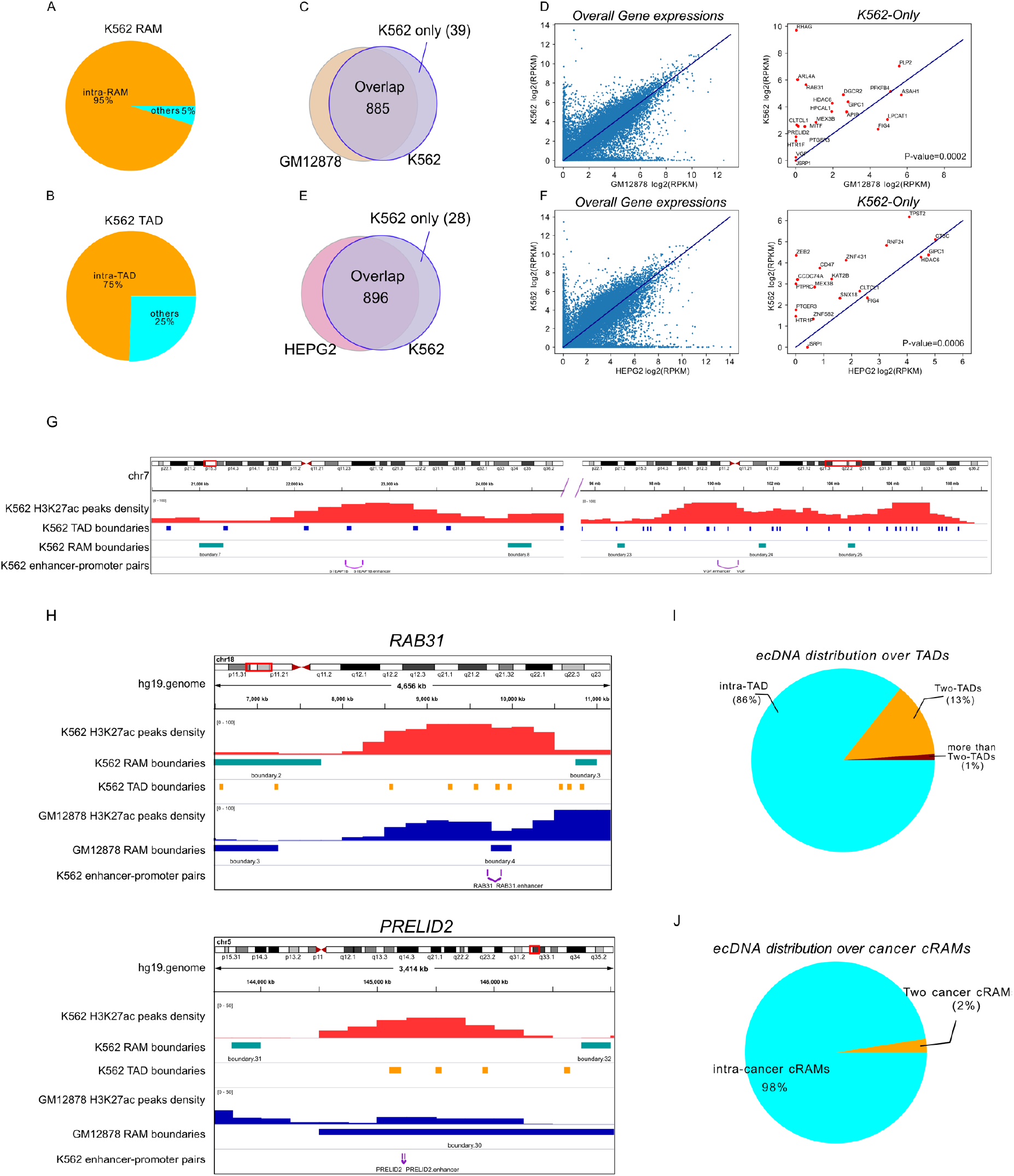
RAM is a functional unit. **(A)** K562 enhancer-promoter pairs distribution over K562 RAMs. **(B)** K562 enhancer-promoter pairs distribution over K562 TADs. **(C)** K562 enhancer-promoter pairs distribution over K562 RAMs and GM12878 RAMs. (**D**) Genes regulated by the 39 enhancer-promoter interactions only within K562 RAMs tend to have higher expressions in K562 compared to GM12878. **(E)** K562 enhancer-promoter pairs distribution over K562 RAMs and HEPG2 RAMs. (**F**) Genes regulated by the 28 enhancer-promoter interactions only within K562 RAMs tend to have higher expressions in K562 compared to HEPG2. (**G**) Examples of K562 enhancer-promoter pairs relative to K562 TAD and RAM boundaries. (**H**) Examples of K562 enhancer-promoter pairs relative to K562 and GM12878 RAM boundaries. (**I**) ecDNA distribution over TADs. (**J**) ecDNA distribution over cancer cRAMs.

We further investigated how the modularity defined by RAMs affects gene expression. To this end, we examined whether the enhancer-promoter pairs located within the same K562 RAMs but separated by RAM boundaries in other cells would specifically impact gene expression in K562. When comparing K562 to the normal cell line GM12878, we found 885 K562 enhancer-promoter interactions were within the same RAMs in both cell lines and 39 only in K562 (**Figure 3C**). The majority of the genes regulated by the 39 enhancer-promoter interactions are upregulated in K562 compared to in GM12878 (P-value=0.0002 by Hypergeometric Test, see **Materials and Methods**) (**Figure 3D**), indicating that the RAM organization facilitates promoter-enhancer interactions to activate gene expression. For example, the *RAB31* promoter interacts with an enhancer that is located within the same RAM in K562 but in a RAM boundary in GM12878 where the enhancer would be silenced in GM12878; the *PRELID2* promoter and its interacting enhancer are located within the same K562 RAM but reside in a GM12878 RAM boundary indicating suppression in GM12878 (**Figure 3H**). In fact, the *RAB31* and *PRELID2* normalized expression levels are 34.29 and 5.37 folds higher in K562 than in GM12878, respectively. All the upregulated gene expressions involved in enhancer-promoter interactions that occurred in the same RAM in K562 but in different GM12878 RAMs are shown in **Table S2**. We had a similar observation by comparing K562 and HEPG2: while the majority of the promoter-enhancer pairs are intra-RAM in both cell lines, 28 are only occurred in the same RAM in K562 but in different HEPG2 RAMs (**Figure 3E**) and the corresponding genes have higher expressions in K562 than in HEPG2 (P-value=0.0006 by Hypergeometric Test) (**Figure 3F, Table S3**). Furthermore, we examined the K562 enhancer-promoter pairs in K562-specific RAMs and the cancer consensus RAMs (cancer cRAMs). 750 pairs were identified as intra-RAM interactions in both K562 RAMs and cancer cRAMs (**fig S4A**). 77 genes were involved in the 174 pairs that are only intra-RAM interactions in K562, and 20 out of the 77 genes were detected as K562 specifically highly expressed genes across 92 cancer cell lines (hypergeometric test P-value=0.03) documented in the Harmonizome database(*27*) (**fig S4B**). These observations further illustrated that RAMs represent a modularity directly associated with functional activity of the chromatin.

### RAMs are insensitive to cohesin degradation

Previous studies showed that cohesin degradation would disrupt loop domains and topological associated domains (TADs) but largely not change histone modifications and gene expression(*28*, *29*). Therefore, RAMs are not expected to be affected by cohesin degradation. For confirmation, we identified RAMs using H3K27ac data in the HCT-116 RAD21-mAC cells untreated and treated for 6 hours with IAA. The RAM patterns for each chromosome were highly correlated between treated and untreated cells (**fig S5A**) and the recall rate for the RAM boundaries was 0.9 on average for all the chromosomes (**fig S5B**). This observation indicates that the RAM formation is independent from cohesin, distinguishing RAMs from TADs and loop domains.

### Extrachromosomal DNA (ecDNA) from cancer patients majorly originated from intra-RAM

Circular extrachromosomal DNAs (ecDNAs) are prevalent in tumors and their length ranges from 100kbp to megabases, and the genes encoded in the ecDNAs are often amplified in cancers(*30*–*32*). We reason that, if RAMs are functional modules, ecDNAs would form within RAMs because RAM boundaries are highly condensed DNAs that would restrain the transcription of genes residing in ecDNAs. To test this hypothesis, we downloaded the ecDNAs identified from cancer patients(*33*), and filtered the ecDNAs corresponding to the median size of the cancer cRAM length (2.5Mb), i.e. only ecDNAs with size <2.5Mb were kept (i.e. 78% of all the ecDNAs). We found that 98% of 2459 ecDNAs were located within the individual RAMs. As a comparison, we performed the same analysis on TADs. We took the conserved TADs defined in the Dixon et al. study(*4*) and only kept the ecDNAs shorter than 880kbp (68% of all the ecDNAs), the median size of the TAD length. We found that 86% of the 2150 ecDNAs were within individual TADs (**Figure 3J, 3K**). Because the ecDNAs were filtered to have comparable length with the RAM and TAD sizes, respectively, this lower intra-domain percentage for TAD compared to RAM is not due to the larger size of RAMs. Furthermore, GREAT analysis(*34*) (http://great.stanford.edu/public/html/) on the ecDNAs that fall into intra-cancer cRAMs but split in TADs revealed that they are highly enriched in “positive regulation of DNA replication” with a P-value of 5.1641E-19. There are 12 genes involved in this pathway: *ATF1*, *BMP5*, *BMP6*, *EGFR*, *FGFR1*, *GLI2*, *IGF1*, *IL6*, *JUN*, *KITLG*, *PDGFA*, *PDGFRA*, which are known important for cell proliferation and cancer pathogenesis. For example, *EGFR* is a driver of tumorigenesis(*35*). Deregulation of the oncogenic FGFR signaling has been frequently observed in multiple types of cancers(*36*). The PDGF mediated signaling has been reported to be involved in the cell proliferation and invasion(*37*). The observations that ecDNAs tend to originate from intra-RAMs, which suggests that RAM is a functional module.

### Deletion of the cRAM boundaries are predicted to alter the 3D chromatin structures

To systematically examine the impact of deleting cRAM boundaries to the chromatin structure, we resorted to computational predictions using a deep learning model ORCA(*38*) (https://github.com/jzhoulab/orca) as it is prohibitive to perform hundreds of Hi-C experiments with sufficient resolution. We took the ORCA model pre-trained on the high resolution Hi-C and Micro-C data in H1-hESC and HFF cell lines to predict 3D chromatin architecture from kilobase to whole-chromosome scale using DNA sequences. It also provided perturbation predictions if certain sequences were targeted. cRAM boundaries shared between cancer and normal samples are apparently important, therefore we selected all 418 of them that are located at least 16Mb away from centromere to predict their impacts on Hi-C contacts if deleted. As a comparison, we also included 298 H1-hESC and 187 HFF TAD boundaries (length of the TAD boundaries >= 100kbp; nonoverlap with selected cRAM boundaries; away from centromere at least 16Mb) in the computational perturbations screening. Considering that the cRAM boundaries are often larger than the TAD boundaries, we only deleted the center 100kbp of cRAM and TAD boundaries to avoid bias introduced by deletion size.

To measure the similarity between the deletion and wildtype Hi-C contact matrices, we calculated the Pearson correlation between them. Compared to the TAD deletions, deleting cRAM boundaries obviously resulted in lower correlation coefficients, indicating larger chromatin alterations, at the highest resolutions the ORCA model could predict (4kb and 8kb resolutions with Wilcoxon Rank Sum test P-values of 2.4E-11 and 2.6E-7, respectively) (Figure 4B). Deletion of the cRAM boundary (*chr10:115,940,000-116,040,000*, in hg38) on Hi-C contacts in HFF and H1-hESC cells is shown as an example (**Figure 4A, fig S6**). The 3D contacts are severely weakened by deleting the cRAM boundary in both cell types.

**Figure 4.**
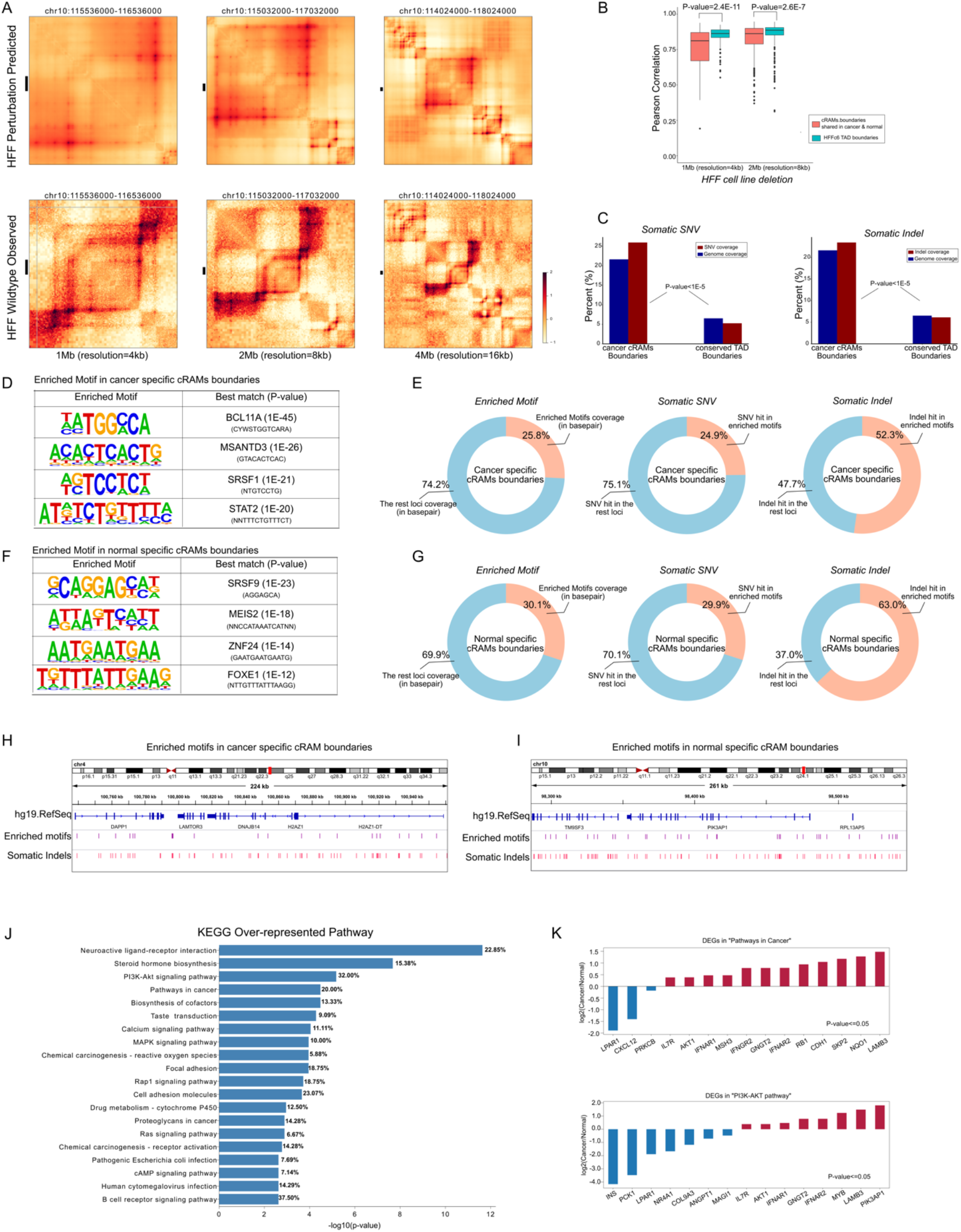
The association of the cRAM boundaries with the 3D chromatin structure and cancer somatic variants. **(A)** An example of the Hi-C contact change upon deletion of the cRAM boundary (chr10:115,940,000-116,040,000 in hg38) in HFF cells predicted by a deep learning model ORCA. (B) Pearson correlations between the predicted Hi-C contacts before and after cRAM boundary and TAD boundary deletion in HFF cells. A lower correlation indicates a larger perturbation to the wildtype chromatin structure upon deletion. (**C**) Somatic SNV and indels enrichment in cancer cRAM boundaries and TAD boundaries. Genome coverage: the total base pairs of the cancer cRAM boundaries or TAD boundaries in the whole genome; SNV coverage: the percentage of the SNVs in the cancer cRAM boundaries or TAD boundaries in the whole genome; INDEL coverage: the percentage of the indels in cancer cRAM boundaries or TAD boundaries in the whole genome. (**D**) Examples of the enriched motifs in the cancer specific cRAM boundaries. (**E**) Overlaps of somatic SNVs and indels with the enriched motifs in the cancer specific cRAM boundaries (**F**) Examples of the enriched motifs in the normal specific cRAM boundaries. (**G**) Overlaps of somatic SNVs and indels with the enriched motifs in the normal specific cRAM boundaries (**H**) The zoom-in genomic view for the enriched motifs with somatic indels in cancer specific cRAM boundaries (**I**) The zoom-in genomic view for the enriched motifs with somatic indels in normal specific cRAM boundaries (**J**) KEGG over-represented pathways for the genes within 2.5kb from the enriched motifs overlapping with somatic indels (**K**) Differentially expressed genes in KEGG “Pathways in cancer” and “PI3K-AKT pathway”

### Somatic genetic variations enriched in regulation associated modules boundaries

If RAMs are functional modules important for regulating functional activities, we reason that somatic mutations in cancers may target the RAM boundaries to disrupt the modular organization of chromatin leading to aberrant regulation of gene expression and resulting tumorigenesis. The PCAWG study revealed consensus mutations and variations from thousands of cancer patients including ~20 millions of somatic single nucleotide variations (SNVs) and ~1.08 millions of indels (https://dcc.icgc.org/releases)(*39*). We found that, while cancer cRAM boundaries cover ~21.6% of the genome, they host 25.9% of somatic SNVs and 23.4% somatic indels. As a comparison, the conserved TAD boundaries(*4*) covering 6.5% genome containing 5.2% somatic SNVs and 6% somatic indels (**Figure 4C**). The cancer cRAM boundaries are significantly enriched with both somatic SNVs and indels compared to the TAD boundaries (P-value<1E-5 from Two-sample Proportion tests, see **Materials and Methods**), indicating a stronger association with cancer mutations.

To elucidate the sequence features associated with cRAM boundaries and investigate how the somatic mutations change such features, we performed motif analysis on the cRAM boundaries using Homer(*40*). We focused on the cRAMs that are common in cancer (normal) but not in normal (cancer) samples as cancer (normal)-specific cRAMs, as they represent changed modularity between cancer and normal samples. By comparing cancer-specific and normal-specific cRAMs, we found 73 and 74 motifs enriched only in cancer and normal specific cRAM boundaries, respectively (example motifs shown in **Figure 4D, 4F**). We employed FIMO(*41*) to identify the occurrences of the enriched motifs that counted for 25.8% and 30.1% in base pairs, respectively, in the cancer and normal specific cRAM boundaries (**Figure 4E, 4G**). We next mapped the PCAWG somatic SNVs and indels onto the cancer and normal specific cRAM boundaries. While somatic SNVs do not show a preferred occurrences within the enriched motifs (24.9% and 29.9% for cancer and normal specific cRAM boundaries, respectively), the somatic indels overlapping with the cancer/normal-specific cRAM boundaries preferentially hit the enriched motifs (52.3% and 63% for cancer and normal specific cRAM boundaries, respectively), more than two-fold by chance, in the altered cRAM boundaries between normal and cancer samples. We speculate that the enriched motifs in the cancer and normal-specific cRAM boundaries may respectively facilitate disruption and formation of cRAM boundaries in the normal samples. Two examples of these motifs overlapping with indels are shown in **Figure 4H-I**. We identified genes within 2.5kb from the enriched motifs overlapping with somatic indels and analyzed the enriched pathways using g:Profiler(*42*). The KEGG over-represented pathways are shown in **Figure 4J,** and the gene ontology molecular functions are shown in **fig S7**. Furthermore, we downloaded the normalized gene expressions of the TCGA and GTEx samples from Expression Atlas (https://www.ebi.ac.uk/gxa/home) and identified differentially expressed genes (DEGs) (P-value<=0.05 by the Wilcoxon Rank Sum tests between cancer and normal samples). The top ranked pathways associated with cancer and cell proliferation are highly enriched with the DEGs, such as *PIK3AP1*, *LABM3*, *AKT1*, *MYB* in the “PI3K-AKT pathway” (32%) and “pathways in cancer” (20%) (**Figure 4K**). These observations suggested that the motifs specifically enriched in the formation of cancer cRAM boundaries and disruption of normal cRAM boundaries are close to genes important for tumorigenesis, cell survival and cell proliferation. Somatic indels can severely alter these motifs and may contribute to the cRAM boundary change, affecting the expressions of the nearby genes.

## DISCUSSION

In this study, we analyzed the peak density profiles of histone modifications data and found they show modular patterns. These modules are clearly defined by active marks such as H3K27ac, H3K4me1 and H3K4me3, indicating their association with functional activity of the genome, and are thus termed regulation associated modules (RAMs). While TADs and compartments are identified from the 3D contacts measured by Hi-C, RAMs are delineated by histone modifications that are directly related to chromatin accessibility and gene expression. We showed that RAMs are obviously distinct from TADs, compartments and LADs although some RAM boundaries do overlap with TAD, compartment or LAD boundaries.

By surveying 93 normal and 19 cancer samples, we found the following evidence to support that RAMs are spatial modules resulting from functional activities. First, we observed that on average 60% of the RAMs (i.e. consensus RAMs) are largely shared across samples, while some of them are sample specific. Compared to TADs, consensus RAMs host higher percent of experimentally confirmed promoter-enhancer contacts (i.e. within the same RAMs), suggesting RAMs represent a modularization of the genome at a scale better aligned with transcriptional regulation. Second, ecDNAs detected from cancer patients tend to originate from the same RAMs rather than across multiple RAMs, supporting the insulation effect of RAM boundaries. Third, deletion of the cRAM boundaries would result in more severe chromatin alteration than the TAD boundaries based on in silico predictions of Hi-C contacts, suggesting the importance of cRAM boundaries in maintaining the chromatin structure. Fourth, cRAM boundaries are also more enriched with somatic genetic variants of SNVs and indels than the TAD boundaries. In particular, the somatic indels tend to disrupt the motifs specifically enriched in cancer or normal specific cRAM boundaries, suggesting a possible mechanism of tumorigenesis involved in altering the chromatin modularity.

To investigate the mechanisms underlying the RAM formation, we found that the RAMs are separated by densely packed DNA regions (as shown by their large number of Hi-C contacts) enriched with repressive histone modifications and lacking open chromatin, active histone marks or transcriptional events. Furthermore, unlike TADs, RAMs are insensitive to cohesin degradation. Taken together, these observations clearly show that RAMs are distinct from loop domains and TADs. RAMs are also different from lamina associated domains (LADs) defined by measuring the intermediate filament protein *LMNB1* localization. The LADs are formed through interactions between chromatin and lamina, and they are located at the periphery of the genome. The RAM boundaries are demarcated by densely packed DNAs and many RAM boundaries are not located in regions interacting with lamina or overlapping with TAD boundaries, and thus the mechanism underlying these RAM boundary formation should be different from other chromatin modules including TADs, LADs and compartments.

Many studies (such as in ref (*43*, *44*)) have shown that multivalent cations such as calcium, magnesium, and manganese can reduce the electrostatic repulsions between the DNA chains and induce DNA condensation. Furthermore, these cations may bind to specific DNA sequences(*45*) and affect nucleosome positioning(*46*, *47*). Therefore, a possible mechanism can be that genomic DNAs become densely packed around cations such as Ca^2+^, Mg^2+^ and Mn^2+^ to form RAM boundaries even if they are not marked by H3K9me3 or interacting with lamina. Proteins such as calcium binding proteins that carry many cations or their interacting partners may recognize specific DNA sequences such as those motifs enriched in the cancer or normal specific cRAM boundaries to facilitate locus-specific localization of cations. Interestingly, the most enriched molecular function of the genes close to (<2.5kbp) the enriched motifs overlapping with somatic indels in the cancer or normal specific cRAM boundaries is calcium ion binding (**fig S7**), and ~30% of them are differentially expressed in cancer and normal samples (**Supplementary Table S4**), implying a possible feedback mechanism. This hypothesis and the mechanistic details are awaiting for future studies.

## Acknowledgements

We are grateful to the labs of Drs. Bas van Steensel, Erez Lieberman Aiden and Marion Cremer for data sharing.

## Funding

This project was partially supported by NIH (R01HG009626 to W.W.).

## Author Contributions

W.W. conceived and supervised the project. L.Z. performed the bioinformatics analysis. L.Z. and W.W. wrote the manuscripts.

## Competing interests

The authors declare no competing interests.

## Data availability

All data needed to evaluate the conclusions in the paper are present in the paper and/or the Supplementary Materials.

## Materials and Methods

### Regulatory associated modules (RAM) identification

#### Data Source

The 93 normal and 19 cancer samples with the called H3K27ac narrow peaks in hg19 were downloaded from Roadmap Epigenomics portal (https://egg2.wustl.edu/roadmap/web_portal/)( *19*) and ENCODE portal (https://www.encodeproject.org/). Table S1 lists the samples used in this study.

#### RAM identification in individual samples

We calculated the H3K27ac narrow-peak density using a sliding window of step size equal to 10kb, 50kb, 100kb, 250kb and 500kb respectively and 500kb flanking size of each window in every sample. The H3K27ac narrow-peak profiles were then smoothed by a local polynomial regression fitting(*48*). The RAM boundaries (valley/minima on the smoothing curves) and peaks (summit/maxima on the smoothing curve) in the smoothed curves were then detected using the “findpeaks” function in R package “pracma”.

#### Consensus-RAM (cRAM) identification

We first identified RAM boundaries in the 93 normal or 19 cancer samples using different step sizes (10kb, 50kb, 100kb, 250kb and 500kb) and then counted the percentage of a genomic region identified as RAM boundary in the 93 normal or 19 cancer samples. A genomic region with occurring percentage >=25% was considered as a consensus-RAM (cRAM) boundary in the normal or cancer samples. We merged cRAM boundaries if they are located <250kb apart from each other and required cRAMs have size >250kb.

### Hi-C data analysis

#### Hi-C processing

The Hi-C data for the wildtype K562, GM12878, A549, IMR90, NHEK, HUVEC, HMEC and the HCT116 cell lines were downloaded from GEO (GSE63525) and the ENCODE portal (ENCSR662QKG and GSE104333). All the raw fastq files were aligned to hg19 genome and then processed using Juicer with the default settings (*7*, *49*). The contact reads in a given cell line were further normalized by vanilla coverage (VC) normalization using the Juicer pipeline. The significance for a given fragment contact was computed by Poisson distribution with VC-normalized expected contact reads versus the VC-normalized observed contact reads. We then used HiCExplorer(*50*–*52*) and HiCplotter(*53*) software to visualize the Hi-C data.

#### A/B compartment

We performed A/B compartment analysis at 250kb resolution. The eigenvectors for each chromosome in all the cell lines involved in the Hi-C data analysis were extracted from the VC normalized Hi-C counts processed by the Juicer pipeline with the default parameters(*54*). POLR2A ChIP-seq data were obtained from the ENCODE(*55*) portal (https://www.encodeproject.org/). To determine A or B compartment, we calculated the correlation between the first eigenvector of each chromosome and the Pol II peak density(*56*). As there was no Pol II ChIP-seq data available for HMEC, we used TSS density in the hg19 genome to assign A/B compartments in HMEC.

#### Topological associated domains (TAD)

To identify topological associated domains (TAD), we applied the insulation score method(*57*) to the Hi-C data at 50kb resolution in the K562, GM12878, A549, IMR90, NHEK, HUVEC, HMEC and HCT116 cell lines. The HFF cell line TADs were downloaded from the 4DN website (accession number 4DNFIMROE6N4). The H1 cell line TADs were downloaded from ref (*4*).

### Lamina-associated domains (LADs) data processing

The Lamin-B1 signal data in the K562, HCT116, H1, HAP1, RPE-hTERT, HFFc6 cell lines generated by the DamID technique were obtained from the 4DN portal(*58*). We downloaded the mean of the replicates for each cell line. The Lamin-B1 signals at 50kb resolution for the K562, HCT116, H1, HAP1, RPE-hTERT, HFFc6 cell lines were lifted over from hg38 to hg19.

### Cohesin degradation analysis

The H3K27ac ChIP-seq data in the untreated HCT-116 RAD21-mAC cells and HCT-116 RAD21-mAC cells treated for 6 hours with IAA were downloaded from GEO (GSE104888). We processed the H3K27ac data same as ref (*59*). In brief, we aligned the raw data to the hg19 human genome using the BWA software(*60*), and then deduplicated the reads using PicardTools. The narrow peaks were called by comparing the associated input data using MACS2(*61*). All the parameters were set to the defaults.

### Enrichment of upregulated genes in enhancer-promoter pairs occurring in the same RAM of K562 but split in the background cell type by Hypergeometric Test

The hypergeometric test was employed to measure the significance of the upregulated genes involved in the enhancer-promoter pairs occurring in the same RAM in K562 (foreground cell type) but in different RAMs of the background cell type. The population size N was the overall genes involved in the K562 and the compared cell type. The population success size M was the number of all upregulated genes in the K562 compared to the background cell type. The sampling size n was the number of the genes involved in the enhancer-promoter pairs occurring in the same RAM of K562 but in different RAMs of the background cell type, and the sampling success size m was the upregulated genes involved in the enhancer-promoter pairs occurring in the same RAM of K562 but in different RAMs of the background cell type. Enrichment was considered significant if p-value<0.05.

### Enrichment of the somatic variants in cancer cRAM boundaries compared to the TAD boundaries assessed by Two-sample Proportion Tests

The two-sample proportion test null hypothesis was to test the equal proportion of the number of the somatic variants relative to the genome coverage (in base pair) in the cancer cRAM boundaries and TAD boundaries. The two proportions were calculated separately by the number of the somatic variants divided by the boundary length (in base pair) for cancer cRAM boundaries and TAD boundaries. Enrichment was considered significant if p-value<0.05.

### Somatic mutations and structural variation analysis for cancer patients from PCAWG

The consensus somatic SNV and indels were downloaded from PCAWG. (https://dcc.icgc.org/releases/PCAWG)(*62*, *63*). The VCF files were transformed to bed files by BEDOPS vcf2bed tools (https://bedops.readthedocs.io/en/latest/content/reference/file-management/conversion/vcf2bed.html)(*64*). The number of somatic SNV and indels overlapping with the RAM and TAD boundaries were then counted.

### Motif analysis

The motif analysis was done using the Homer pipeline (*40*) with default parameters. The motif occurrence was called using FIMO (*41*) with p-value <=1 E-4.

## Notes

### Competing Interest Statement

The authors have declared no competing interest.

